# AURORA: a longitudinal single-cell framework for resolving autophagy-associated remodeling under nutrient stress

**DOI:** 10.1101/2025.03.12.642827

**Authors:** Joel D. Posligua-Garcia, M. C. Banqueri-Pegalajar, Carlos Ulises Cárdenas-Vela, Anabel Martínez-Padilla, James R. Perkins, C. Rodriguez-Caso, Jose Luis Urdiales, Juan A. G. Ranea, Miguel Ángel Medina, Manuel Bernal

**Affiliations:** Department of Molecular Biology and Biochemistry, Faculty of Sciences, University of Malaga, Andalucía Tech, 29071 Malaga, Spain; Biomedical Research Institute of Malaga and Nanomedicine Platform, IBIMA BIONAND Platform, 29590 Malaga, Spain; CIBER de Enfermedades Raras (CIBERER, Spanish Network of Research Center in Rare Diseases), Instituto de Salud Carlos III, E-28029 Madrid, Spain; Spanish National Bioinformatics Institute (INB/ELIXIR-ES), Instituto de Salud Carlos III, E-28029 Madrid, Spain

**Author notes:** Corresponding authors: Miguel Ángel Medina and Manuel Bernal.

**Keywords:** Autophagy, nutrient deprivation, live-cell imaging, high-content imaging, single-cell heterogeneity, longitudinal cell tracking, phenotypic states

## Abstract

Nutrient stress induces dynamic and heterogeneous cellular responses that are often reduced to population averages or endpoint measurements. AURORA is a longitudinal single-cell phenotyping framework integrating high-content time-lapse imaging, trajectory-aware quality control, multi-compartment morphology, reference phenotypic states, and cross-cell-line aggregation of state-occupancy behaviors.

AURORA is applied to MCF-7, MDA-MB-231, A-549, and HT-1080 cancer cells during 4 h exposure to Earle’s balanced salt solution (EBSS). Population-level Total Autophagy shows sustained EBSS-associated elevation in MCF-7, A-549, and HT-1080, but a stronger early response followed by attenuation in MDA-MB-231. Longitudinal analysis shows that these averages arise from cell-line-and time-dependent changes in single-cell distributions rather than uniform displacement. Feature-level analysis further identifies distinct combinations of nuclear, whole-cell, and autophagosome-associated organization, compactness, and shape in each model.

Projection of EBSS-treated cells into Complete-defined phenotypic landscapes reveals line-specific redistribution among reference states. Despite this specificity, dominant state-occupancy signatures converge into six recurrent higher-order programs, linking different cellular routes to partially shared remodeling architectures. Because DAPGreen is not paired with a flux inhibitor or ratiometric reporter, these findings describe an autophagy-associated imaging phenotype rather than autophagic flux. AURORA therefore distinguishes population-average change from heterogeneous single-cell adaptation during dynamic stress responses.

## Introduction

Autophagy is a central adaptive pathway that recycles intracellular material and reorganizes metabolic resources during stress. In cancer, however, its consequences are strongly context-dependent: autophagy can limit damage, but it can also support tumor-cell survival, plasticity, and metabolic fitness under fluctuating environmental constraints [1–5]. Nutrient deprivation is therefore widely used to probe autophagy-associated stress responses, although the resulting behavior depends on both the cellular background and the specific nutrient context [6–8].

Most autophagy responses are still summarized using bulk or endpoint measurements. These readouts establish whether a condition changes an average signal, but they cannot determine whether the response is rapid or delayed, sustained or transient, or uniformly shared across cells. This distinction is especially relevant in cancer, where phenotypically distinct cells can coexist and respond differently to the same stress [9]. Population averages may consequently conceal redistribution between lower-and higher-responding subpopulations as well as divergent trajectories followed by individual cells.

Live-cell imaging provides the temporal ordering and longitudinal identity needed to resolve this heterogeneity [10,11]. Nevertheless, autophagy-focused imaging often centers on puncta or intensity measurements that require careful interpretation and do not, by themselves, establish autophagic flux [12,13]. Nutrient stress can also remodel cell shape, nuclear organization, and the spatial arrangement of intracellular compartments [14]. Integrating these features with single-cell trajectories can therefore describe stress adaptation as a dynamic phenotypic process rather than as a single reporter value. Previous live-cell phenotyping frameworks have demonstrated the value of trajectory-aware state analysis for capturing cellular behaviors that are inaccessible to static snapshots [15–18].

Here, we developed AURORA, a longitudinal single-cell phenotyping framework that links population-level dynamics, single-cell distributions, multi-compartment morphology, and phenotypic state occupancy within the same live-cell experiment. We applied AURORA to MCF-7, MDA-MB-231, A-549, and HT-1080 cancer cells exposed to Earle’s balanced salt solution (EBSS), a commonly used nutrient-deprivation medium [19], during a 4 h imaging window. Rather than seeking a universal starvation phenotype, we asked whether responses that appear similar at the population level arise from shared or cell-line-specific temporal and single-cell remodeling, and whether distinct line-specific states can be organized into recurrent higher-order phenotypic programs.

## Results

### AURORA distinguishes population-level responses from integrated longitudinal single-cell remodeling

We first established the AURORA workflow to determine how much information is gained when a starvation response is examined beyond population averages (Fig. 1). Cells maintained in Complete medium or exposed to EBSS were imaged every 20 min for 4 h, generating 13 timepoints per field after a standardized staining and equilibration sequence (Fig. 1A). This common experimental design allowed the same image series to be analyzed at two distinct but complementary biological levels.

**Figure 1.**
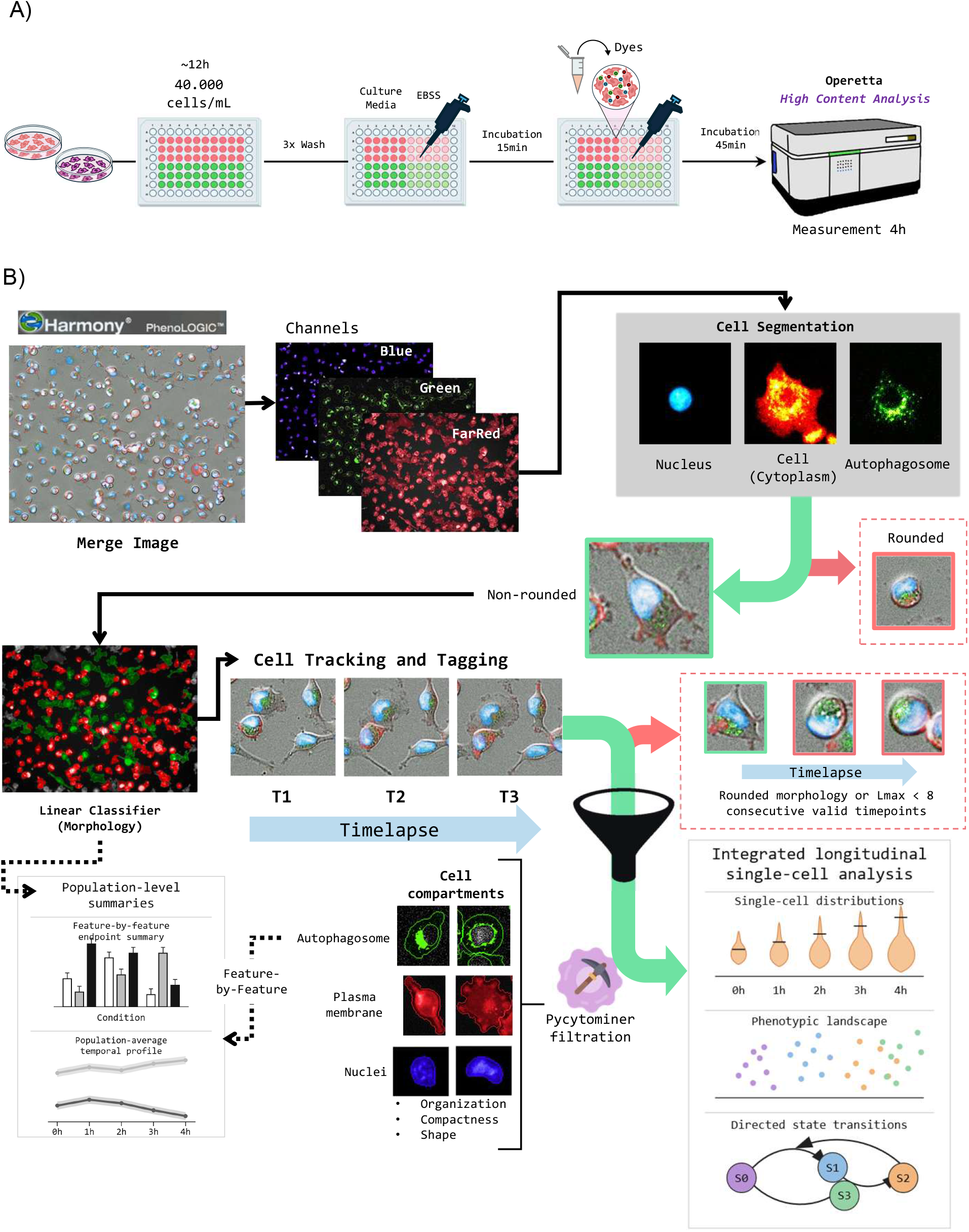
Experimental workflow and single-cell image-analysis pipeline. (A) Live-cell imaging workflow. MCF-7, MDA-MB-231, A-549 and HT-1080 cells are plated at 40,000 cells/mL and allowed to attach (∼12 h) before a 3× wash into complete growth medium or EBSS starvation medium, a 15 min incubation, addition of fluorescent reporter dyes, a 45 min dye-loading incubation, and continuous imaging on an Operetta High-Content Analysis system over a 4 h time course. (B) Image-analysis pipeline. Multichannel images (Blue/Green/FarRed) are segmented in Harmony/PhenoLOGIC into nucleus, cell (cytoplasm) and autophagosome compartments. Black arrows trace the main processing workflow; green and red arrows mark the morphology-based filters applied along it (green = retained, red = excluded — first rounded vs. non-rounded cells, then tracks with a rounded morphology at any timepoint or fewer than 8 consecutive valid timepoints). Dashed arrows lead to the population-level outputs, where each morphological feature is summarized and analyzed one at a time (feature-by-feature endpoint and temporal summaries); solid arrows lead instead to the integrated longitudinal single-cell analyses detailed in Figures 2–5, where all extracted, PyCytominer-filtered features are combined together for each cell (single-cell distributions, phenotypic-landscape projections, directed state-transition models).

At the population level, measurements from the segmented cells were reduced to condition-and timepoint-level summaries (dashed branch in Fig. 1B). This branch quantified Total Autophagy and evaluated individual morphological descriptors separately, producing endpoint comparisons and population-average temporal trajectories. It therefore captured whether the average signal increased, decreased, or changed over time, but it did not preserve the identity of individual cells or reveal whether a mean response arose uniformly across the population.

The integrated single-cell branch retained longitudinal identity after nuclear, whole-cell, and DAPGreen-positive autophagosome-associated segmentation (solid branch in Fig. 1B). Compartment-resolved fluorescence, size, shape, compactness, and spatial-organization descriptors were linked across successive images for each tracked cell. Rounded cells, border-associated objects, and discontinuous trajectories were excluded. Minimum-consecutive-track evaluation identified k = 8 as the operational continuity threshold (Supplementary Fig. 1A). Replicate-ordering diagnostics supported the consistency of population-level measurements (Supplementary Fig. 1B), while final valid-track counts documented the analyzable cohort retained for each cell line and condition (Supplementary Fig. 1C). Net-displacement distributions were included as a final longitudinal tracking QC measure (Supplementary Fig. 1D).

Retaining cell identity exposed biological structure that population means cannot resolve. The single-cell branch quantified the full distributions of cellular responses, projected EBSS-treated cells onto phenotypic states defined exclusively from Complete-condition cells, and reconstructed state occupancy and directed transitions over time (Fig. 1B). These state-level behaviors were subsequently aggregated into recurrent cross-cell-line programs. Figure 1 therefore defines the central analytical comparison used throughout the Results: population-level analysis describes the average magnitude and kinetics of starvation-associated change, whereas longitudinal single-cell analysis determines whether that change reflects uniform behavior or heterogeneous redistribution across cells, states, and trajectories.

### Longitudinal AURORA profiling captures cell-line-dependent autophagy-associated dynamics

We first examined population-average Total Autophagy over the 4 h recording. EBSS increased the signal relative to Complete medium in all four cell lines, but the magnitude and temporal persistence of the response differed markedly among models (Fig. 2A,B). At the 4 h endpoint, the EBSS-associated increase was significant in MCF-7, A-549, and HT-1080, whereas MDA-MB-231 was not significantly different from Complete (Fig. 2A).

**Figure 2.**
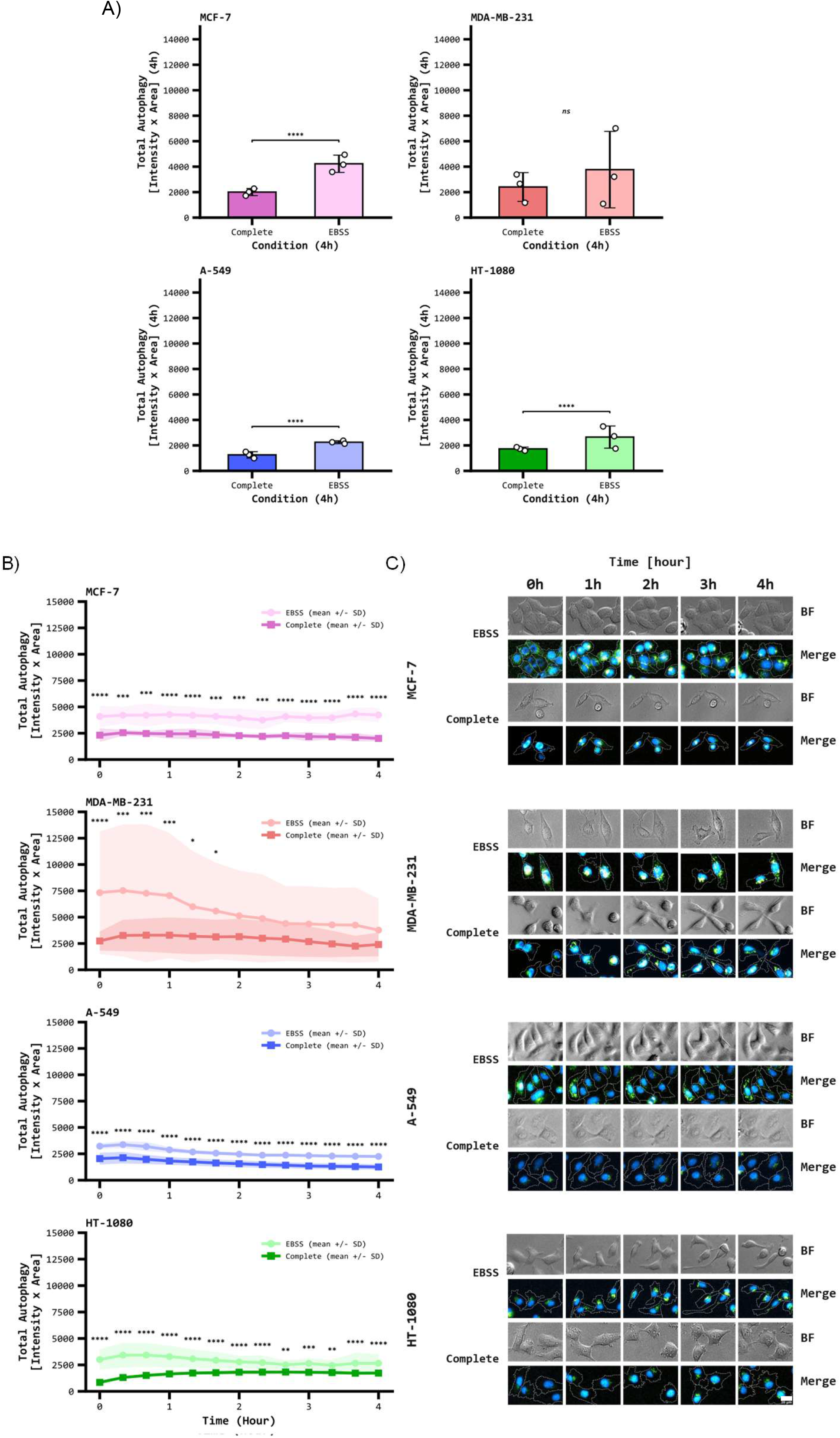
Population-level dynamics of EBSS-induced autophagy signal. Total Autophagy signal (Complete vs. EBSS) for MCF-7, MDA-MB-231, A-549 and HT-1080. (A) Endpoint comparison at 4 h (timepoint 13). Bars show mean ± SD (n = 3 biological replicates); points are individual replicate means; asterisks denote statistical significance (ns = not significant; see Statistics below). (B) Full 4 h time course (13 timepoints, 20 min intervals); asterisks denote significance at each timepoint (Statistics below). (C) Representative brightfield (BF) and merged-fluorescence (Merge) time-lapse images (0–4 h) of the same field for one Complete- and one EBSS-treated well per cell line, tracked across the full time course; scale bar = 20 µm. *Statistics (A–B):* paired t-test (Complete vs. EBSS) using the pooled residual error from a two-way repeated-measures ANOVA (Time × Condition; Subject = Replicate). (A) reports this test as a single isolated comparison at 4 h (uncorrected); (B) reports the same test at each of the 13 timepoints, Bonferroni-corrected across the timepoint family (*p < 0.05, **p < 0.01, ***p < 0.001, ****p < 0.0001).

MCF-7 maintained a clear EBSS-versus-Complete separation throughout the time course, with EBSS values remaining approximately twofold higher despite a modest decline in both conditions (Fig. 2B). A-549 and HT-1080 also displayed sustained separation across the 13 timepoints. Their absolute Total Autophagy values were lower than those of MCF-7 and MDA-MB-231, emphasizing that the effect of EBSS should be interpreted within each cell line rather than through direct comparison of raw signal magnitude between lines.

MDA-MB-231 showed a distinct transient profile. Total Autophagy was highest under EBSS at the beginning of the experiment and progressively decreased, while the Complete trajectory remained comparatively stable. Accordingly, timepoint-level differences were concentrated in the early phase and were no longer detected during the later timepoints, consistent with the nonsignificant 4 h endpoint comparison (Fig. 2A,B).

Representative time-lapse images were concordant with these quantitative trajectories (Fig. 2C). EBSS-treated MCF-7, A-549, and HT-1080 cells retained visibly stronger DAPGreen-associated signal across the imaging window, whereas the contrast between conditions diminished over time in MDA-MB-231. Replicate-resolved global feature-dispersion scores showed that the condition-dependent broadening was observed across biological replicates (Supplementary Fig. 2A), and the pooled distributions summarized its direction and magnitude within each line (Supplementary Fig. 2B). Cell counts remained broadly stable during the 4 h experiment (Supplementary Fig. 2C). In the separate prolonged assay, SYTOX-positive nuclei accumulated under EBSS in a strongly cell-line-dependent manner, particularly after the early 4 h imaging window (Supplementary Fig. 2D).

Thus, acute nutrient deprivation did not elicit a single population-level response: it produced sustained autophagy-associated remodeling in MCF-7, A-549, and HT-1080 but a comparatively transient response in MDA-MB-231. These differences motivated analysis of the single-cell distributions underlying the population means.

### Single-cell longitudinal distributions reveal heterogeneous autophagy-associated remodeling

Split-violin distributions revealed broad Total Autophagy heterogeneity in every cell line and condition throughout the experiment (Fig. 3A). Complete and EBSS populations overlapped, but EBSS altered their median, spread, and upper tails in a cell-line-and time-dependent manner. The population-average differences in Fig. 2 therefore did not reflect synchronized displacement of all cells.

**Figure 3.**
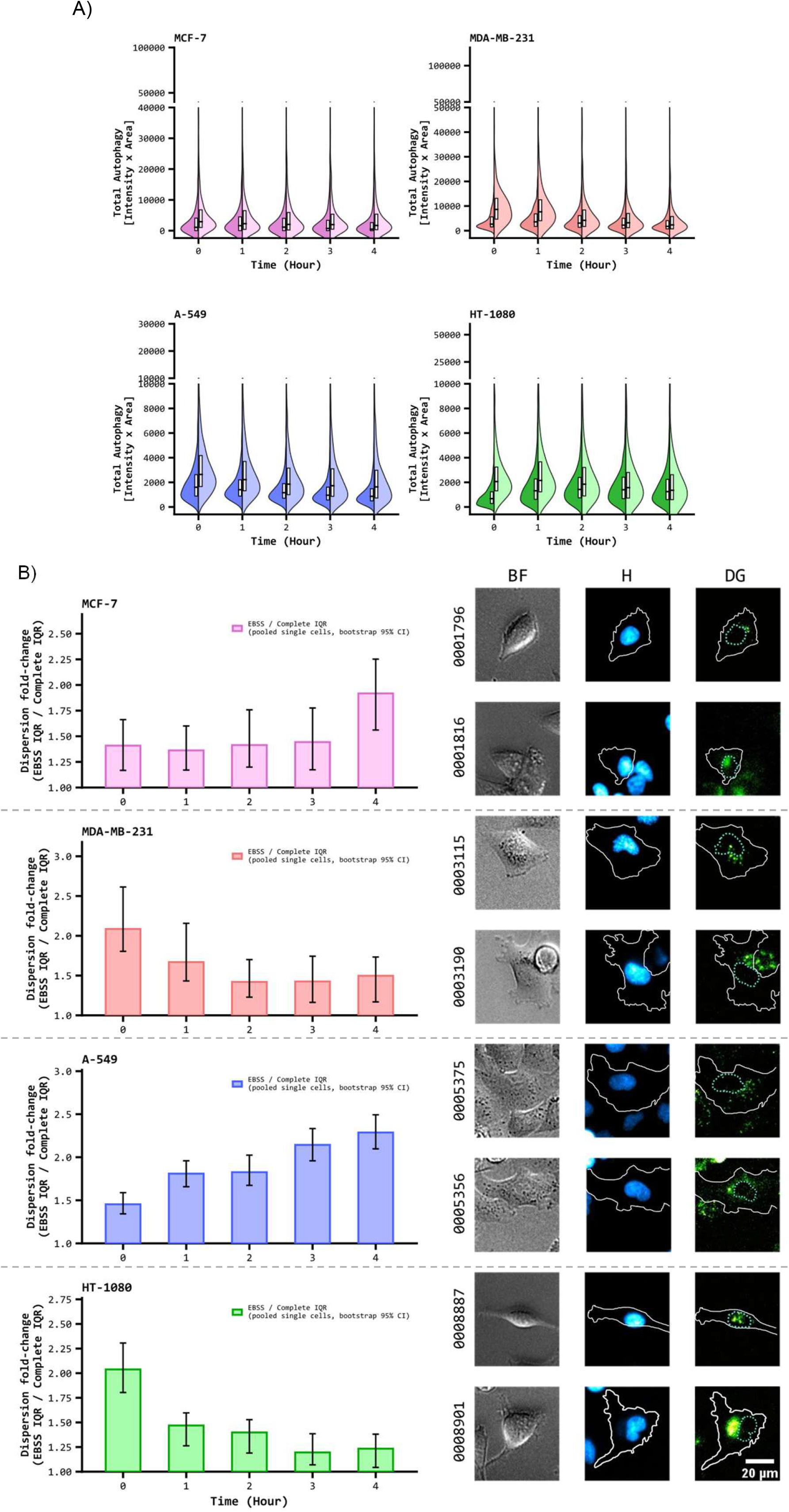
Single-cell dispersion of EBSS-induced autophagy remodeling. Single-cell dispersion of Total Autophagy across the 4 h time course, Complete vs. EBSS. (A) Split-violin distributions (Complete, left half; EBSS, right half) at each timepoint, one panel per cell line; box shows interquartile range and median. (B) Dispersion fold-change (EBSS IQR / Complete IQR) of pooled single-cell values over time, one panel per cell line; error bars are bootstrap 95% CI (2,000 resamples). Representative brightfield (BF), Hoechst (H) and autophagosome-dye (DG) images show two EBSS-treated cells from the same field at the timepoint of peak dispersion for that cell line (MCF-7 and A-549: 4 h; MDA-MB-231 and HT-1080: 0 h; see panel B). Within each cell line, the upper row is the lower-Total-Autophagy example and the lower row is the higher-Total-Autophagy example. The number printed to the left of each row is that cell’s longitudinal Track ID; scale bar = 20 µm. All panels use the quality-controlled single-cell track cohort (≥8 consecutive valid timepoints per track; see Fig. 1B).

Dispersion fold-change, calculated as the EBSS-to-Complete interquartile-range ratio, remained above 1 at every displayed timepoint in all four cell lines (Fig. 3B). MCF-7 showed a moderate early increase in dispersion that became largest at 4 h. A-549 exhibited a progressive rise, reaching its greatest dispersion at 4 h. In contrast, MDA-MB-231 and HT-1080 displayed their largest fold-changes at the first timepoint, followed by attenuation, although dispersion remained elevated relative to Complete.

The timing of maximal dispersion was therefore not shared across models: it occurred at 4 h in MCF-7 and A-549 but at the first imaging timepoint in MDA-MB-231 and HT-1080 (Fig. 3B). For each cell line, the representative EBSS-treated cells are arranged from lower Total Autophagy signal at the top to higher signal at the bottom. The number printed to the left of each image row is the longitudinal Track ID of that cell, allowing the displayed example to be linked unambiguously to its tracked single-cell trajectory. These paired examples illustrate the range of DAPGreen-associated signal observed at the line-specific timepoint of maximal dispersion.

These observations establish that similar changes in population-average Total Autophagy can emerge from different distributional architectures. Acute starvation reshaped the relative abundance of low-and high-signal cells rather than driving each population uniformly, providing a rationale for examining the broader morphological descriptors associated with this heterogeneity.

### Feature-level analysis identifies cell-line-specific remodeling architectures

Starting from the common 83-feature descriptor panel, pycytominer retained 65 features for MCF-7, 65 for MDA-MB-231, 58 for A-549, and 56 for HT-1080 (Fig. 4A). Nuclear, whole-cell, and autophagosome-associated measurements were represented in every model, with autophagosome-associated descriptors forming the largest compartmental fraction in A-549 and a similarly prominent fraction in the other lines. Forty-four features were shared by all four selected sets, while the remaining intersections demonstrated partial line-specific feature usage (Fig. 4B). The engineered feature-group definitions are provided first in Supplementary Table 1, and the complete feature-usage matrix then identifies which descriptors were retained for each cell line and how every descriptor maps onto the shared interpretative axes (Supplementary Table 2).

**Figure 4.**
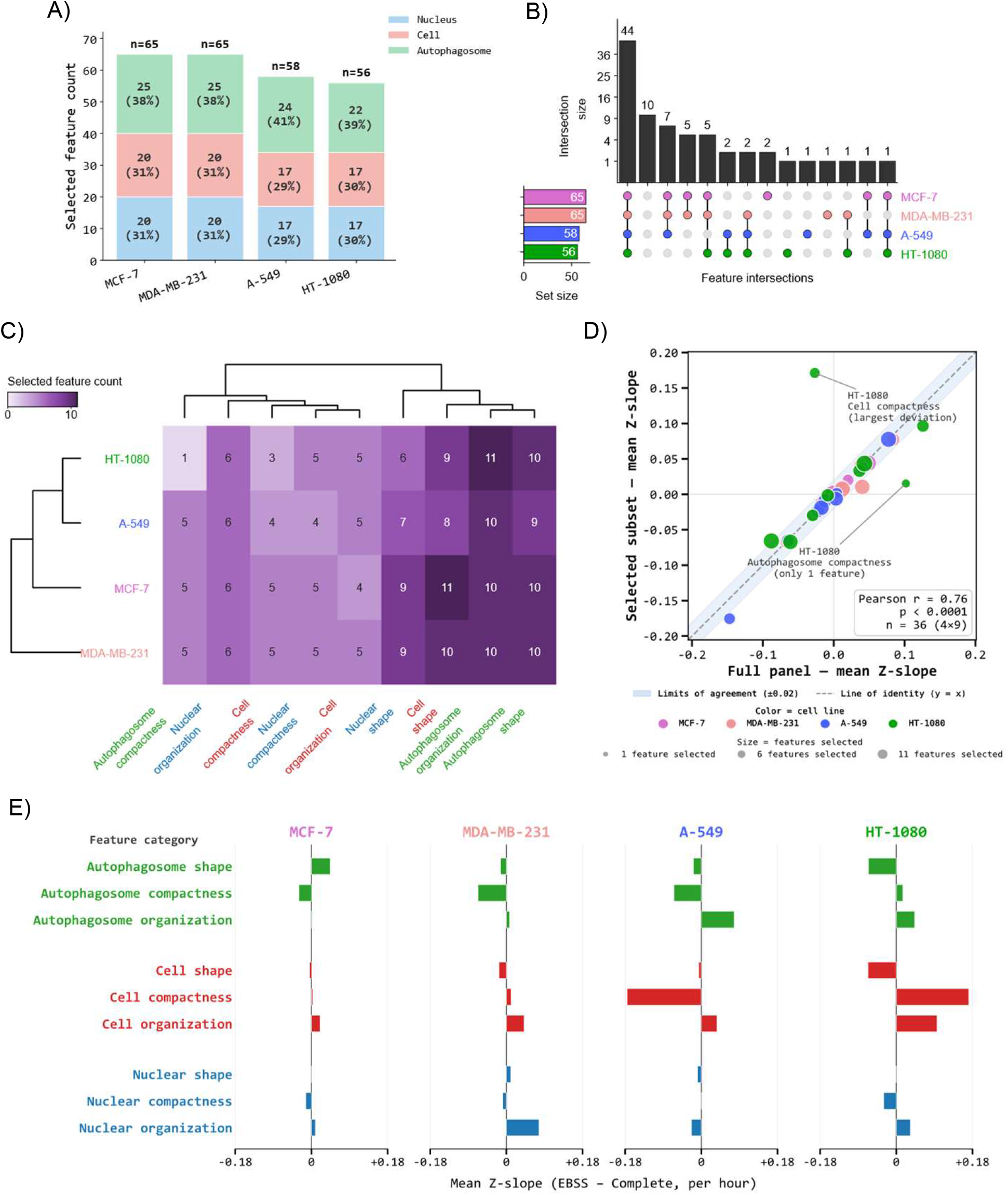
Feature architecture and EBSS-response concordance of the pycytominer-selected morphological feature set. (A) Number of pycytominer-selected features per cellular compartment (Nucleus, Cell, Autophagosome), for each cell line. (B) UpSet plot of selected-feature overlap across the four cell lines. (C) Hierarchically clustered heatmap of the nine meta-feature groups (organization / compactness / shape × Nucleus / Cell / Autophagosome) across the selected features. (D) Concordance between EBSS-response slopes computed on the full feature set vs. the pycytominer-selected subset. Each point is one meta-feature category × cell line (n = 36); slope = weighted-least-squares linear trend of the EBSS-minus-Complete gap over elapsed time, from the pooled residual error of a two-way repeated-measures ANOVA design (Subject = Replicate × Timepoint × Condition). Pearson r = 0.76, p = 7.3×10^⁻^ (n = 36). Shaded band: ±0.02 limits of agreement around the identity line; the two largest deviations (HT-1080 Cell compactness; HT-1080 Autophagosome compactness) are labeled. (E) EBSS-response slopes (selected feature set) by meta-feature category, one tornado chart per cell line; bars colored by compartment (Nucleus, Cell, Autophagosome). Neither panel uses significance testing; both report point estimates (slopes) only.

Because the selected feature sets differed among cell lines, direct comparison of individual model features would mix biological differences with unequal feature availability. We therefore summarized the descriptors into nine meta-features formed by crossing three compartments (nucleus, whole cell, and autophagosome-associated objects) with three property classes (organization, compactness, and shape). These shared axes reduce correlated measurements to biologically interpretable categories while preserving the cell-line-specific selected features for state-model fitting. The clustered category counts demonstrate how strongly each shared axis was represented in each model (Fig. 4C). Definitions of the engineered feature groups are provided in Supplementary Table 1, and the descriptor-to-meta-feature mapping and cell-line-specific selection membership are provided in Supplementary Table 2.

We next tested whether the selected features preserved the temporal EBSS-response architecture observed with the full descriptor panel. Across the 36 cell-line-by-meta-feature combinations, EBSS-minus-Complete temporal slopes from the selected subsets correlated with those from the full panel (Pearson r = 0.76, p = 7.3 × 10⁻□; Fig. 4D). Most estimates lay close to the identity line and within the ±0.02 agreement band; the largest deviations involved HT-1080 whole-cell compactness and autophagosome compactness. This concordance shows that the meta-feature summaries remain representative of the broader descriptor response despite model-specific feature selection.

The selected-feature slopes nevertheless exposed distinct remodeling signatures (Fig. 4E). MCF-7 combined increased autophagosome shape with reduced autophagosome compactness; MDA-MB-231 was distinguished by decreased whole-cell compactness and increased nuclear organization; A-549 combined reduced whole-cell compactness with increased autophagosome organization; and HT-1080 showed increased whole-cell compactness and organization together with reduced autophagosome shape. Therefore, EBSS engaged different compartmental axes in each cell line rather than producing a universal morphological feature shift.

### Complete-defined state projection reveals cell-line-specific phenotypic redistribution

To determine how these feature changes reorganized cellular phenotypes, we defined cell-line-specific states from Complete-condition cells and projected EBSS cells into the corresponding reference landscapes (Fig. 5A). For each line, the left UMAP shows the Complete-condition landscape used to define the reference states and the right UMAP shows EBSS cells projected into that same embedding. The fixed state contours and labels therefore provide a common within-line coordinate system rather than separate clustering of the two conditions. MCF-7 and MDA-MB-231 each resolved three states, A-549 resolved four, and HT-1080 resolved three. Changes in density within and across the fixed contours indicated that EBSS redistributed cells through the pre-existing Complete-defined landscapes.

**Figure 5.**
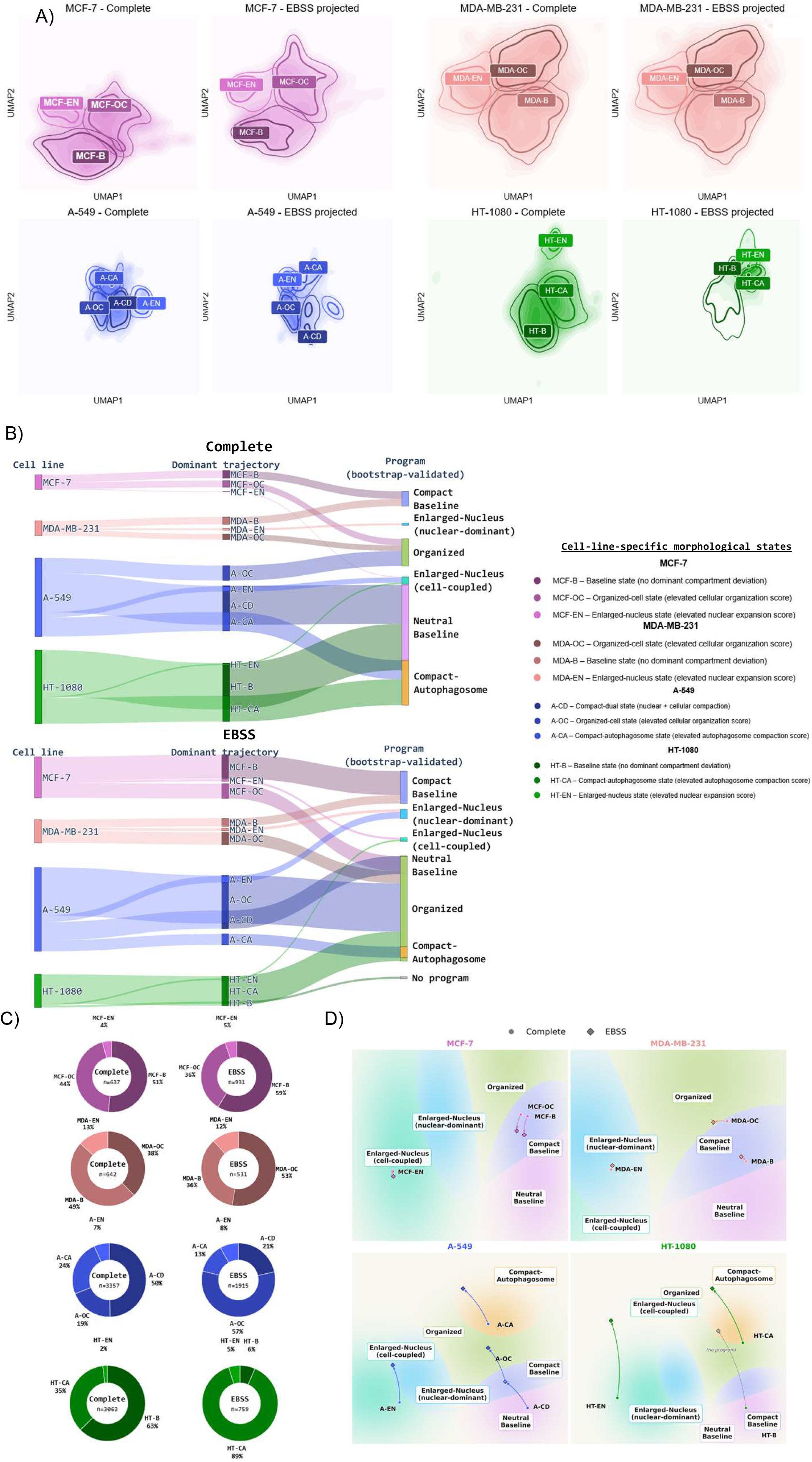
Cell-line-specific morphological states. Per-cell morphological states, defined independently for each cell line by K-means clustering (fit on Complete-condition cells only, on robust z-scored pycytominer-selected features) and projected onto EBSS cells with an elastic-net logistic classifier trained to recover the Complete-defined states. (A) Per-cell UMAP state landscapes (Complete vs. EBSS), one per cell line; UMAP fit on Complete cells and EBSS cells transformed into the same embedding. (B) Sankey diagrams (cell line → dominant trajectory → cross-cell-line trajectory program), one per condition. Programs are a bootstrap co-clustering consensus (1,000 resamples; HDBSCAN on 1 − co-clustering probability) yielding six cross-cell-line programs plus a residual “No program” group. (C) Donut charts of dominant-trajectory occupancy (one vote per cell, most frequent observed state), Complete vs. EBSS, one per cell line. (D) Waddington-style potential landscape of the shared program space, shown as one combined panel per cell line (2×2 layout). All four panels are cropped views of the exact same two-dimensional projection and the same bootstrap-validated program regions from (B) — only the zoom window differs, centered on each cell line’s own data footprint; a program not visible or labeled in a given panel is therefore not absent in principle, that cell line’s cells simply do not populate that region of the shared landscape within the panel’s cropped view. Arrows show each state’s centroid displacement from Complete (circle) to EBSS (diamond).

The Sankey representation then connected each cell line to its dominant state trajectories and from those trajectories to the recurrent cross-cell-line programs (Fig. 5B). Displaying Complete and EBSS separately showed that the same line-specific state could contribute to a shared higher-order program while the relative flow through states and programs changed after starvation. This panel establishes the correspondence between cell-line-specific labels and the program-level terminology used below.

Dominant-trajectory occupancy quantified the underlying state redistribution with one vote per retained track (Fig. 5C). In MCF-7, the baseline state increased from 51% under Complete to 59% under EBSS, while the organized-cell state decreased from 44% to 36%. MDA-MB-231 shifted in the opposite organizational direction: its organized-cell state increased from 38% to 53%, whereas the baseline state decreased from 49% to 36%. The enlarged-nucleus fractions changed only modestly in both breast-cancer lines.

The largest dominant-state redistributions occurred in A-549 and HT-1080 (Fig. 5C). In A-549, the organized-cell state increased from 19% to 57%, accompanied by reductions in the compact-dual state from 50% to 21% and the compact-autophagosome state from 24% to 13%. In HT-1080, the compact-autophagosome state expanded from 35% to 89%, while the baseline state contracted from 63% to 6%; the enlarged-nucleus state increased from 2% to 5%.

### Higher-order aggregation identifies recurrent phenotypic programs across cell lines

Bootstrap co-clustering of the 26 CellLine × Condition × State groups identified six supported programs—Organized, Compact-Autophagosome, Enlarged-Nucleus (nuclear-dominant), Enlarged-Nucleus (cell-coupled), Compact Baseline, and Neutral Baseline—plus one residual No program group (Fig. 5B). Mapping the dominant trajectories onto these programs showed that the same higher-order configuration could be reached through different line-specific states. The clearest EBSS-associated changes were the expansion of Organized through A-549’s organized-cell state and of Compact-Autophagosome through HT-1080’s compact-autophagosome state (Fig. 5B,C).

The common two-dimensional program landscape completed the ordered main-figure analysis (Fig. 5D). Complete and EBSS centroids for each state were displayed in the same coordinate system and against the same program regions, although each cell-line panel used a cropped view centered on its own data footprint. The arrows therefore summarize state-specific displacement between conditions without implying that an unlabeled program is absent from the shared landscape.

The supplementary state diagnostics support this interpretation in panel order. Meta-feature heatmaps first define the Complete and EBSS signatures of every line-specific state (Supplementary Fig. 3A). Dominant-state donut charts then provide the condition-specific occupancy summaries corresponding to Fig. 5C (Supplementary Fig. 3B). Finally, Markov diagrams visualize the descriptive transition topology, state occupancy, and self-transitions for each cell line and condition (Supplementary Fig. 3C).

The final robustness series documents the complete state-to-program analysis in order. The computational schematic summarizes Complete-state definition, EBSS projection, dominant behavior, and bootstrap program assignment (Supplementary Fig. 4A). Complete-state recovery was 99.64-99.81% across lines (Supplementary Fig. 4B). Track-level bootstrap perturbation of the transition matrices produced condition- and line-dependent mean absolute changes, ranging from 0.37 in A-549 Complete to 0.94 in MDA-MB-231 EBSS, supporting descriptive rather than inferential use of the transition diagrams (Supplementary Fig. 4C). Predicted state-occupancy distributions independently recapitulated the principal condition shifts (Supplementary Fig. 4D). Mean within-program co-clustering probabilities were approximately 0.94-0.99 for all six supported programs (Supplementary Fig. 4E), and occupancy-weighted meta-feature profiles confirmed that they reflected distinct combinations of nuclear, whole-cell, and autophagosome-associated organization, compactness, and shape (Supplementary Fig. 4F). Together, these results show how population-level changes resolve into heterogeneous, line-specific state redistribution and partially convergent higher-order programs.

## Discussion

The principal contribution of AURORA is to show that acute nutrient deprivation cannot be adequately described by a single population-average autophagy-associated value. By combining live-cell imaging, longitudinal identity, multi-compartment morphology, and reference-state analysis, the framework resolved four complementary dimensions of the response: its magnitude, temporal persistence, single-cell distribution, and phenotypic organization. Across the four cancer cell lines, EBSS did not produce one stereotyped response. Similar population-level increases could instead arise from different temporal profiles and from redistribution among phenotypically distinct cells (Figs. 2-5).

The first distinction was kinetic. MCF-7, A-549, and HT-1080 maintained an EBSS-associated elevation across the imaging window, whereas MDA-MB-231 showed a stronger early response that progressively attenuated (Fig. 2). The divergence was particularly informative in the two breast cancer models. Previous EBSS experiments comparing these same lines reported an earlier LC3B-II maximum in MDA-MB-231 than in MCF-7, together with distinct transcriptional responses among autophagy-related and signaling genes [19]. Although the readouts and experimental windows differ, both studies support the conclusion that starvation-associated dynamics are strongly conditioned by cellular background. More generally, time-resolved analyses have shown that different stages of the autophagy pathway can respond with distinct onset and persistence and that the nutrient removed influences the resulting kinetics [7,8]. Because EBSS simultaneously removes several classes of nutrients and growth factors, the present data do not identify the upstream sensor responsible for each trajectory. They do show, however, that a 4 h endpoint alone would collapse sustained and transient responses into the same type of comparison. For acute stress phenotyping, response shape is therefore biologically informative in addition to endpoint magnitude.

The second distinction was distributional. Complete and EBSS populations remained partially overlapping even when their means diverged, and the degree and timing of dispersion differed among cell lines (Fig. 3). Thus, the population response reflected changing proportions of lower- and higher-signal cells rather than synchronous movement of every cell. Earlier live-cell studies also identified substantial cell-to-cell variability in starvation-associated autophagy dynamics [7,8], and recent work in pancreatic cancer demonstrated that local extracellular-matrix sensing can generate spatially heterogeneous autophagic flux with functionally different cellular consequences [20]. AURORA extends this general principle in a controlled two-dimensional setting by retaining each cell’s longitudinal identity and placing signal heterogeneity within a broader morphological context. Because DAPGreen was not paired with a flux inhibitor or a ratiometric flux reporter here, this comparison concerns heterogeneity in an autophagy-associated imaging phenotype, not direct heterogeneity in autophagic flux.

The third distinction was architectural. Feature selection did not converge on an identical descriptor set in all four lines, and EBSS engaged different combinations of nuclear, whole-cell, and autophagosome-associated organization, compactness, and shape (Fig. 4; Supplementary Tables 1 and 2). This result cautions against assigning a universal morphological meaning to an increase in autophagy-associated signal. Autophagy is embedded in cell-wide processes that influence metabolism, adhesion, cytoskeletal organization, and survival, and its consequences depend strongly on cellular context [3,21–23]. The meta-feature representation was useful precisely because it separated two analytical levels: cell-line-specific descriptors were retained for fitting each model, whereas shared compartment-by-property axes allowed the resulting remodeling architectures to be compared without pretending that the underlying selected features were identical.

The Complete-defined reference landscapes provided a fourth level of interpretation. Defining states only from Complete-condition cells and then projecting EBSS cells into those fixed landscapes made the perturbation visible as redistribution relative to each line’s own baseline organization (Fig. 5A-C). This design avoids allowing the stress condition to redefine the reference against which its effect is judged. It also complements approaches that embed full morphological trajectories to reveal dynamic cell-state relationships that are inaccessible to population snapshots [15–18]. In AURORA, the central quantity is not the existence of a cluster by itself, but the change in longitudinal occupancy of reproducibly defined reference states.

Cross-line aggregation then revealed partial convergence above this line-specific state structure. Although the four models used different state labels and followed different occupancy changes, their nine-axis signatures grouped into six recurrent higher-order programs plus one residual group (Fig. 5B-D). This suggests that distinct cell lines can reach related morphological configurations through different baseline states and response routes. The program names are therefore intentionally descriptive: they summarize the dominant morphology of co-clustered signatures and do not define molecular cell types, discrete attractors, or conserved mechanistic pathways. Their strongest value is comparative, providing a common vocabulary for patterns that would otherwise remain trapped within separate cell-line models.

HT-1080 illustrates both the biological relevance and the interpretative limits of this framework. It displayed pronounced early phenotypic redistribution and the largest increase in SYTOX-positive nuclei during the separate prolonged EBSS assay (Fig. 4E; Supplementary Fig. 2D). Nutrient-responsive adaptation has previously been demonstrated in HT-1080 cells: disruption of the GCN2-eIF2α-ATF4 axis during amino-acid limitation elicited an initially cytoprotective autophagic response but reduced survival when adaptation failed [24]. In a distinct nutrient context, cystine deprivation caused substantial HT-1080 death within 24 h, while lysosomal catabolism of extracellular protein supplied cysteine and suppressed ferroptosis [25]. These studies do not establish the mechanism operating under EBSS, but they show that HT-1080 survival is strongly conditioned by nutrient composition and lysosome-linked adaptive capacity. Our data therefore motivate the hypothesis that the marked early displacement observed under EBSS may identify reduced tolerance to sustained nutrient deprivation in this model. Establishing that relationship will require linking early trajectories directly to later fate in the same cells and testing the relevant stress-response and cell-death pathways.

A practical strength of the framework is that these state summaries remain connected to the underlying tracks. Minimum-continuity filtering, final track-count reporting, and displacement diagnostics expose sources of uncertainty that are often hidden when dynamic imaging data are reduced directly to state diagrams (Supplementary Fig. 1). This is important because broken or selectively retained trajectories can distort apparent occupancy and transition frequencies [10]. Accordingly, the Markov diagrams in this study are used as descriptive summaries of observed state sequences rather than as estimates of a stationary or causal transition process; the bootstrap sensitivity analysis makes that boundary explicit (Supplementary Figs. 3C and 4C).

Several limitations define the scope of the conclusions. First, DAPGreen intensity and morphology report autophagosome-associated structures but do not measure pathway completion or autophagic flux [6,12,13]. Tandem fluorescent LC3 reporters, flux inhibitors, or pulse-chase reporter-processing assays would be needed to distinguish increased formation from reduced lysosomal clearance [26,27]. Second, EBSS is a compound nutrient-deprivation perturbation, so the cell-line-specific trajectories cannot be assigned to a single nutrient-sensing pathway. Third, the 4 h window captures early remodeling but not the complete adaptation or fate response; conversely, the separate 24 h viability assay cannot retrospectively assign fate to the cells tracked in the early experiment. Fourth, the study includes four established cancer cell lines in two-dimensional culture. The reproducibility and biological meaning of the programs should therefore be tested in additional genetic backgrounds, primary or patient-derived models, and three-dimensional microenvironments. Finally, morphology-derived states are operational representations of phenotype. Cell-cycle status, metabolic state, and unmeasured molecular programs may contribute to their structure, and molecular profiling or perturbational validation will be required before mechanistic identities can be assigned.

Together, these findings support a multilevel view of acute starvation adaptation. Population averages describe whether an autophagy-associated signal changes; longitudinal single-cell distributions reveal how broadly that change is shared; multi-compartment features identify the architecture in which it occurs; and reference-state occupancy shows how the population is reorganized over time. AURORA connects these levels without treating any one of them as a surrogate for autophagic flux or molecular mechanism. Its immediate value is therefore as a hypothesis-generating phenotyping framework: it identifies when apparently similar responses arise through different cellular routes and defines specific state-and trajectory-level phenotypes for subsequent molecular validation.

## Conclusion

AURORA demonstrates that acute nutrient deprivation cannot be adequately described by a single population-average autophagy-associated measurement. By connecting response kinetics, longitudinal single-cell distributions, multi-compartment morphology, and reference-state occupancy, the framework distinguishes sustained from transient responses and resolves the heterogeneous cellular routes underlying similar average changes. Although the four cancer models follow distinct remodeling trajectories, their dominant state-occupancy signatures organize into six recurrent higher-order phenotypic programs. The current DAPGreen readout describes an autophagy-associated imaging phenotype rather than autophagic flux, but AURORA provides a reproducible framework for generating state- and trajectory-level hypotheses that can be tested through molecular perturbation and direct fate-linked validation.

## Materials and Methods

### Cell lines, culture conditions, and experimental setup

Four human cancer cell lines representing distinct tumor backgrounds were used in this study: MCF-7, a luminal breast adenocarcinoma cell line; MDA-MB-231, a triple-negative/basal-like breast adenocarcinoma cell line; A-549, a lung adenocarcinoma cell line; and HT-1080, a fibrosarcoma cell line. Cell lines were obtained from ATCC-derived stocks and identified according to supplier catalog information: MCF-7 (ATCC HTB-22), MDA-MB-231 (ATCC HTB-26), A-549 (ATCC CCL-185), and HT-1080 (ATCC CCL-121). Cells were maintained under standard culture conditions at 37 °C in a humidified atmosphere containing 5% CO₂. MCF-7 and HT-1080 cells were cultured in high-glucose DMEM, whereas A-549 and MDA-MB-231 cells were cultured in RPMI-1640. Both media were supplemented with 10% fetal bovine serum (FBS; South American origin), 2 mM L-glutamine, and 1x penicillin-streptomycin.

For time-lapse experiments, cells were seeded in 96-well microplates (Thermo Scientific Nunc, Cat. 167008; Nunclon Delta-treated clear polystyrene, flat-bottom, with lid) at density of 40,000 cells/mL and allowed to adhere overnight (∼12 h). Acute nutrient deprivation was induced by replacing complete growth medium with unsupplemented Earle’s Balanced Salt Solution (EBSS; Gibco, Cat. 14155063), whereas complete growth medium was used as the control condition, hereafter referred to as complete. Cells were imaged for 4 h to capture early autophagosome-associated single-cell dynamics.

### Sytox Orange viability assay under prolonged EBSS

To assess cell death during prolonged nutrient deprivation, a separate SYTOX Orange assay was analyzed under Complete and EBSS conditions over 24 h. SYTOX Orange Nucleic Acid Stain (0.25 µM; Invitrogen/Thermo Fisher Scientific, S11368) was used together with Hoechst 33342 (0.1 µM; Invitrogen/Thermo Fisher Scientific, H3570). The figure-generating dataset contained two replicate wells per condition for each cell line, sampled at hourly timepoints from 0 to 24 h. Harmony exported the well-level fraction of Hoechst-positive nuclei that were also SYTOX Orange-positive; this fraction was multiplied by 100 and replicate wells were averaged at each hourly timepoint for visualization. Accordingly, this assay is presented as a descriptive extended cell-death measurement rather than as part of the 4 h single-cell tracking analysis.

### Live-cell staining, imaging, and feature extraction

Live-cell staining was subsequently performed without additional wash steps using Hoechst 33342 (0.1 µM; Invitrogen/Thermo Fisher Scientific, H3570), CellMask Deep Red Plasma Membrane Stain (0.25 µM; Invitrogen/Thermo Fisher Scientific, C10046), and DAPGreen Autophagy Detection dye (2.5 µM; Dojindo, D676). DAPGreen is a small-molecule live-cell probe reported to label forming autophagic structures without transfection, to remain fluorescent across the autophagic process, and to correlate strongly with LC3-positive puncta [28]. Plates were then incubated for 45 min at 37 °C and 5% CO₂ before transfer to the Operetta high-content imaging system. Images were acquired on an Operetta high-content imaging system using a 20x long-working-distance objective every 20 min for 4 h, yielding 13 timepoints. The first acquired image was defined as timepoint 1 of the imaging sequence. Because complete medium or EBSS was added before dye equilibration and staining, cells were exposed to the corresponding condition during the 60 min pre-imaging incubation. For each cell line and condition, three technical replicate wells were imaged per biological replicate, with four fields of view acquired per well. Three independent biological replicates were performed.

Quantitative image analysis was performed in Harmony 4.8 using a standardized segmentation and feature-extraction workflow. Nuclei were segmented from Hoechst signal, whole-cell regions from CellMask Deep Red, and cytoplasmic regions were defined as whole-cell masks excluding nuclear masks. DAPGreen-positive autophagosome-associated structures were detected after background correction. Compartment-resolved morphological, morphometric, and fluorescence intensity features were extracted from nuclei, cytoplasm, whole-cell regions, and DAPGreen-positive objects. In this study, Total Autophagy denotes an image-derived quantitative descriptor calculated for each cell from DAPGreen-positive autophagosome-associated objects detected in Harmony. Operationally, it corresponds to the area-weighted DAPGreen-associated signal, calculated by multiplying the corrected DAPGreen fluorescence intensity of detected autophagosome-associated objects by their segmented area, as detailed below. Therefore, Total Autophagy is interpreted throughout the manuscript as an autophagy-associated imaging readout, not as a direct measurement of total biological autophagy or autophagic flux. Segmentation thresholds were optimized separately for each cell line and kept constant between Complete and EBSS conditions within each line. Cells partially captured at image borders were excluded.

### Single-cell tracking and longitudinal filtering

Longitudinal single-cell tracking was performed in Harmony using object tracking across sequential timepoints. Exported measurement tables were organized into tracked single-cell trajectories across replicates, wells, fields, and timepoints while preserving the corresponding image-derived features for each tracked cell. This provided the longitudinal backbone for downstream filtering, state assignment, transition analysis, and state-occupancy summaries.

Longitudinal filtering was then performed using a minimum consecutive track-length criterion. For each trajectory, the longest uninterrupted sequence of valid measurements across the 13 acquired timepoints was computed. Track retention was evaluated across increasing minimum consecutive length thresholds (k = 6-13; Supplementary Fig. 1A). Based on this retention analysis, k = 8 was selected as the operational threshold for downstream analyses, balancing temporal continuity with retention of sufficient analyzable trajectories across cell lines and conditions. Tracks with fewer than eight consecutive valid timepoints were excluded from downstream analyses.

### Feature engineering and feature selection

Harmony-derived measurements were complemented with engineered descriptors capturing cellular size, shape, compartmental organization, and morphological complexity, resulting in a shared panel of 83 features. Redundant and low-information features were removed separately for each cell line using pycytominer on Complete-condition profiles. The resulting fixed feature sets were used for state modeling, whereas the shared descriptor panel was retained for biological interpretation. Detailed feature definitions, filtering parameters, and the complete feature-to-meta-feature mapping are provided in the Supplementary Methods and supplementary tables.

Features were normalized relative to Complete-condition reference populations before downstream analysis. Matched replicate-timepoint normalization was used for feature-level comparisons, whereas state modeling and cross-cell-line program construction used the pooled Complete population within each cell line. For interpretation across models, normalized descriptors were summarized into nine common axes spanning organization, compactness, and shape in nuclear, whole-cell, and autophagosome-associated compartments. Full normalization rules are provided in the Supplementary Methods.

### Phenotypic state modeling and EBSS projection

Cell-line-specific phenotypic states were defined from pycytominer-selected features using only Complete-condition cells after longitudinal quality control [29]. K-means clustering yielded three states for MCF-7, three for MDA-MB-231, four for A-549, and three for HT-1080. These labels represent operational phenotypic reference states rather than discrete molecular identities.

An elastic-net multinomial logistic-regression classifier was trained [30,31] for each cell line to recover the Complete-defined K-means states and was then applied unchanged to EBSS cells. This projected starvation-treated cells into the same basal reference system used for state occupancy, transition summaries, and higher-order program analysis. Model parameters and recovery diagnostics are reported in the Supplementary Methods and Supplementary Fig. 4B.

### Markov transition analysis and dominant trajectories

Within each cell line and condition, state assignments were ordered along retained single-cell tracks. Empirical transition probabilities were calculated from consecutive timepoint pairs, including self-transitions, and were interpreted descriptively [32,33]. Robustness was assessed by track-level bootstrap resampling within each cell line and condition (Supplementary Fig. 4C).

For dominant-state visualizations, each track contributed one vote corresponding to its most frequently occupied state. Net displacement between the first and last valid centroid coordinates was used only as longitudinal QC. UMAP was used for two-dimensional visualization of the fixed state landscapes [34]. Complete methodological definitions and sensitivity analyses are provided in the Supplementary Methods.

### Cross-cell-line program discovery by bootstrap co-clustering

Cross-cell-line programs were inferred from the nine common morphological axes. Each track was summarized separately within every state it occupied, and the resulting CellLine x Condition x State signatures were compared by bootstrap co-clustering. HDBSCAN was applied to the resulting precomputed distance matrix [35]. Across 1,000 resamples, co-clustering probabilities quantified how reproducibly pairs of state groups converged in the same density-supported cluster.

Final clustering of the co-clustering distance matrix yielded six recurrent programs and one residual No program group. The shared landscape in Fig. 5D was generated from the same group signatures in a common two-dimensional PCA space. Complete aggregation rules, UMAP/HDBSCAN settings, bootstrap seeds, confidence definitions, and PCA preprocessing are provided in the Supplementary Methods.

### Statistical analysis and visualization

Unless otherwise stated, analyses were performed independently for each cell line, with Complete and EBSS compared within the same line. Biological replicates were the experimental units for the 4 h population-level analysis; technical wells and fields were not treated as independent replicates.

Population-level Total Autophagy was analyzed using two-way repeated-measures ANOVA with biological replicate matched across condition and time, followed by Bonferroni-adjusted Complete-versus-EBSS comparisons at individual timepoints. Other QC and distributional analyses were descriptive unless a statistical test is explicitly stated. Detailed aggregation rules, error terms, bootstrap procedures, and software parameters are provided in the Supplementary Methods.

Segmentation, tracking, primary measurement extraction, and well-level SYTOX quantification were performed in Harmony 4.8. Representative images were visualized in ImageJ/Fiji [36,37]. Computational analyses were performed in Python [38] using pycytominer [29], NumPy [39], pandas [40], SciPy [41], scikit-learn [42], and umap-learn [34]. Matplotlib [43] and seaborn [44] were used for static visualization, Plotly [45] for Sankey diagrams, and joblib [46] for model serialization.

## Acknowledgments

The authors thank Sara Molina Gil, technician at ECAI NANOIMAGEN (NANBIOSIS), IBIMA Plataforma BIONAND, for valuable assistance with high-content screening imaging and technical support. The authors also acknowledge the cell culture facility at IBIMA Plataforma BIONAND for providing the infrastructure required for this study. Joel D. Posligua-Garcia, M. C. Banqueri-Pegalajar, and Carlos Ulises Cárdenas-Vela are currently enrolled in the Programa de Doctorado en Biología Celular y Molecular (PhD Program in Cellular and Molecular Biology), Facultad de Ciencias (Faculty of Sciences), Universidad de Málaga (University of Malaga), Spain.

## Funding

This work received limited support from research group BIO-267 (Junta de Andalucía). M. Bernal was supported by the A.4 Fellowship (Ayudas para la Incorporación de Doctores), II Plan Propio, Universidad de Málaga. Juan A. G. Ranea acknowledges funding from the Spanish Ministry of Economy and Competitiveness (PID2022-140047OB-C21) and the EURAS project (Horizon Europe, HORIZON-HLTH-2022-DISEASE-06; project ID 101080580).

## Conflict of interest

The authors declare no conflicts of interest.

## Author contributions

Joel D. Posligua-Garcia (J.D.P.-G.): Investigation, Formal analysis, Visualization, and Writing - original draft. M. C. Banqueri-Pegalajar (M.C.B.-P.), Carlos Ulises Cárdenas-Vela (C.U.C.-V.), and Anabel Martínez-Padilla (A.M.-P.): Investigation and Writing - review & editing. James R. Perkins (J.R.P.), C. Rodriguez-Caso (C.R.-C.), Jose Luis Urdiales (J.L.U.), and Juan A. G. Ranea (J.A.G.R.): Writing - review & editing. Miguel Ángel Medina (M.Á.M.) and Manuel Bernal (M.B.): Conceptualization, Methodology, Supervision, Project administration, and Writing - original draft and review & editing. Miguel Ángel Medina and Manuel Bernal serve as corresponding authors.

## Declaration of generative AI and AI-Assisted technologies

OpenAI GPT-5.6 Sol accessed through Codex, and Anthropic Claude Sonnet 5 were used during manuscript preparation and revision to support language editing, structural refinement, formatting, reference organization, and review of computational documentation. All AI-assisted output was critically reviewed, verified, and revised by the authors. The authors take full responsibility for the scientific content, interpretations, and final text.

## Abbreviations

ANOVA: analysis of variance
AURORA: Analysis of Unified Responses and Organization in Reconfiguring Adaptive systems
DAPGreen: DAPGreen Autophagy Detection dye
DMEM: Dulbecco’s Modified Eagle Medium
EBSS: Earle’s Balanced Salt Solution
FBS: fetal bovine serum
HDBSCAN: hierarchical density-based spatial clustering of applications with noise
IQR: interquartile range
LC3: microtubule-associated protein 1 light chain 3
PCA: principal component analysis
PBS: phosphate-buffered saline
QC: quality control
RPMI-1640: Roswell Park Memorial Institute 1640 medium
UMAP: Uniform Manifold Approximation and Projection

---

**Supplementary Figure 1.**
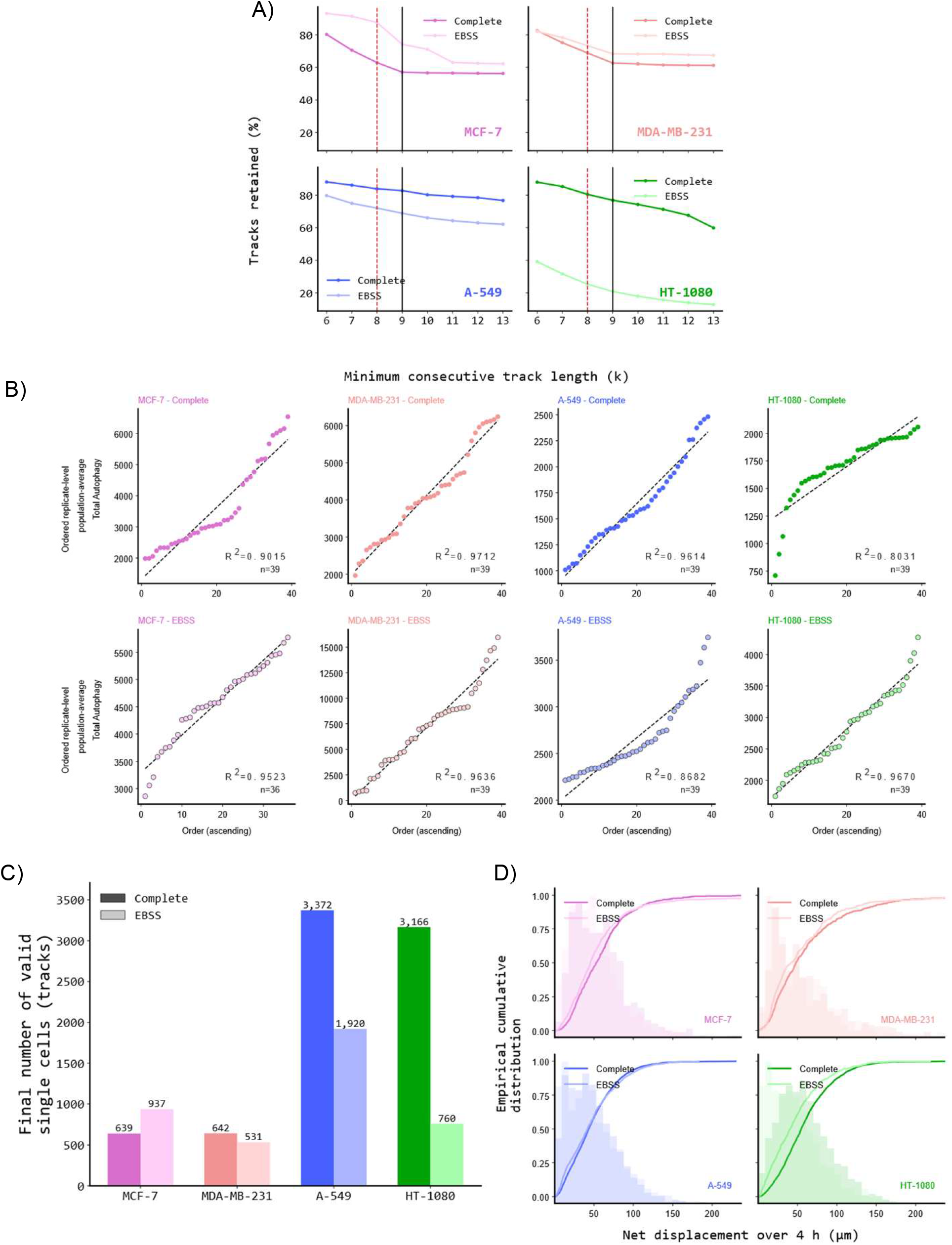
Track-filtering quality control. (A) Percentage of tracks retained at the selected minimum-consecutive-timepoint threshold (k = 8), one bar per cell line × condition (Complete, EBSS). (B) Replicate-level population-average Total Autophagy values, sorted in ascending order and plotted against their rank, one panel per cell line × condition (8 panels); dashed line = linear least-squares fit, R² and n (number of replicate-timepoint values) reported per panel. (C) Final valid single-cell track counts (≥ 2 timepoints with valid X/Y coordinates) per cell line × condition. (D) ECDF (with background density histogram) of net displacement per track over the 4 h recording, Complete vs. EBSS, one panel per cell line; Complete/EBSS difference summarized by Mann–Whitney U.

**Supplementary Figure 2.**
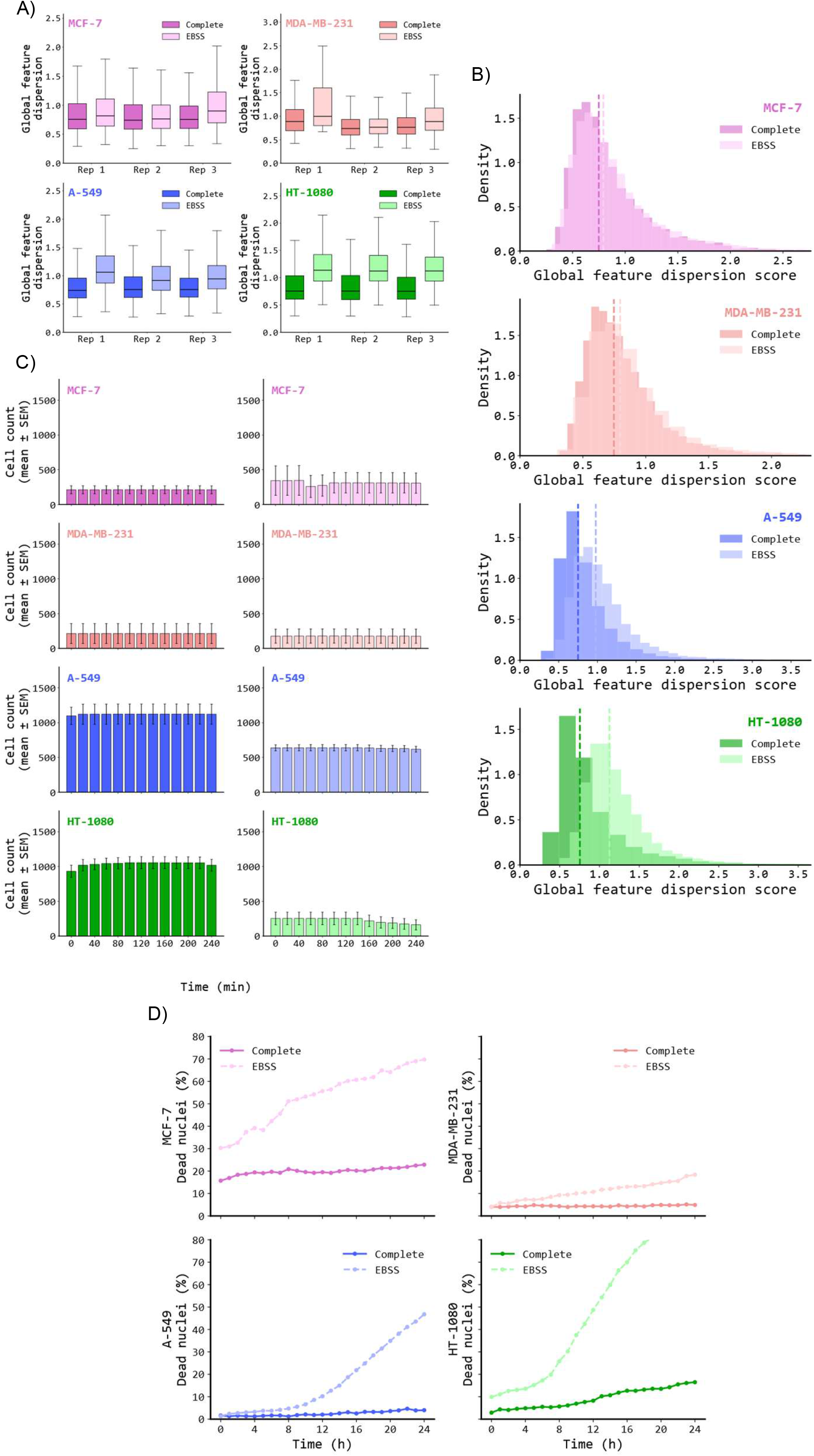
Replicate consistency and cell-count quality control. (A) Global feature-dispersion score (RMS of the z-scored 83-feature morphology vector) of single cells, resolved by replicate, Complete vs. EBSS, one panel per cell line. (B) Pooled density histograms of the same dispersion score per condition, one panel per cell line (x-axis clipped at the 99.5th percentile). (C) Mean single-cell counts per timepoint (mean ± SEM across replicates), one panel per cell line × condition. (D) SYTOX Orange⁺ / Hoechst⁺ nuclei (%) over 24 h, Complete vs. EBSS, independent cell-death assay.

**Supplementary Figure 3.**
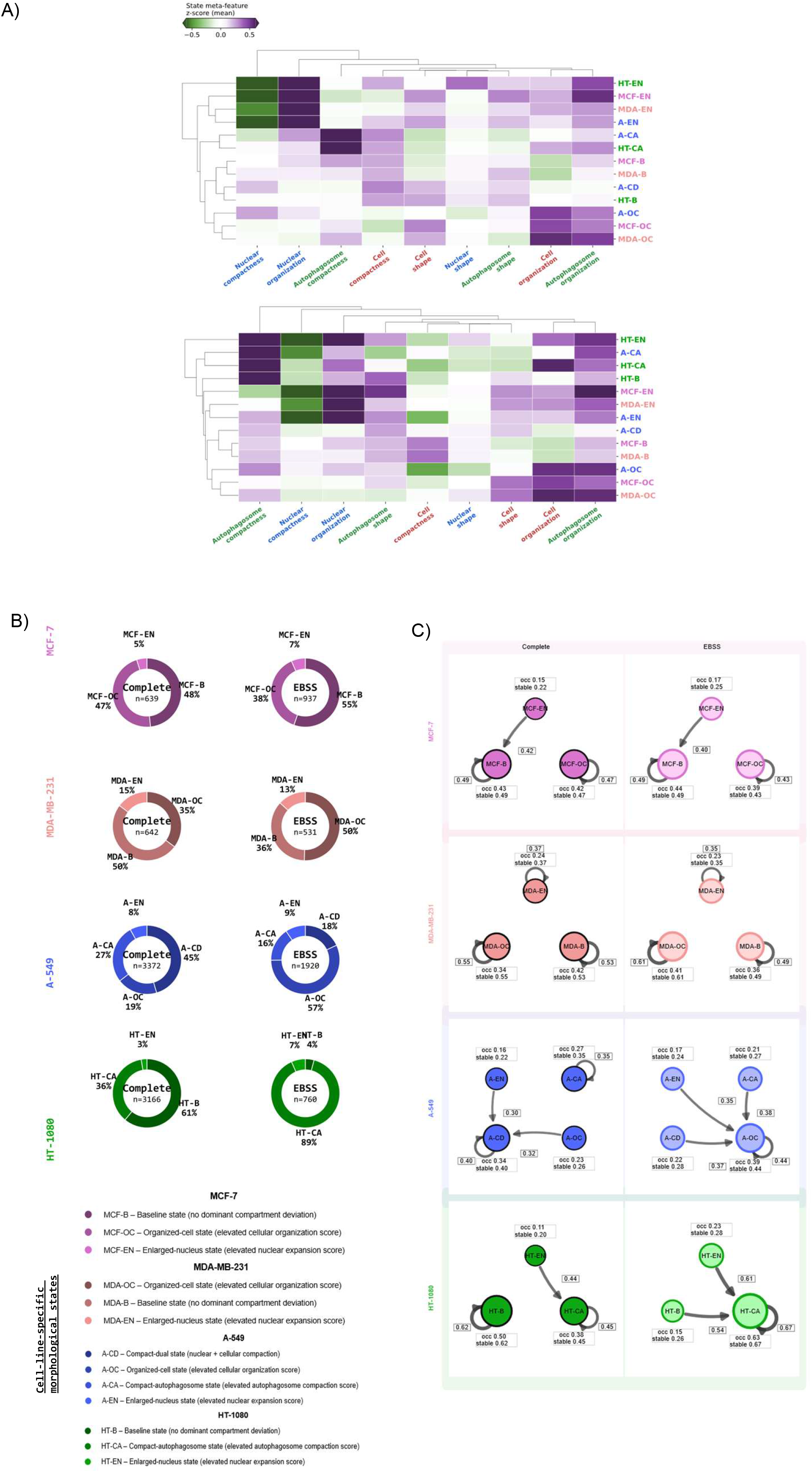
Cell-line-specific morphological states – state-model diagnostics. Diagnostics for the same per-cell state model used in Figure 5, sharing state assignments and the K-means / elastic-net methodology described there. (A) Clustered heatmap of mean meta-feature scores per state, one panel per condition (Complete, EBSS). (B) Donut charts of dominant-trajectory occupancy, Complete vs. EBSS, one per cell line (same design as Figure 5C). (C) Markov state-transition diagrams (bubble diagrams), one per cell line × condition (8 panels, shared axis scale); bubble size = state occupancy, arrow width = transition probability, self-loops shown as return arcs.

**Supplementary Figure 4.**
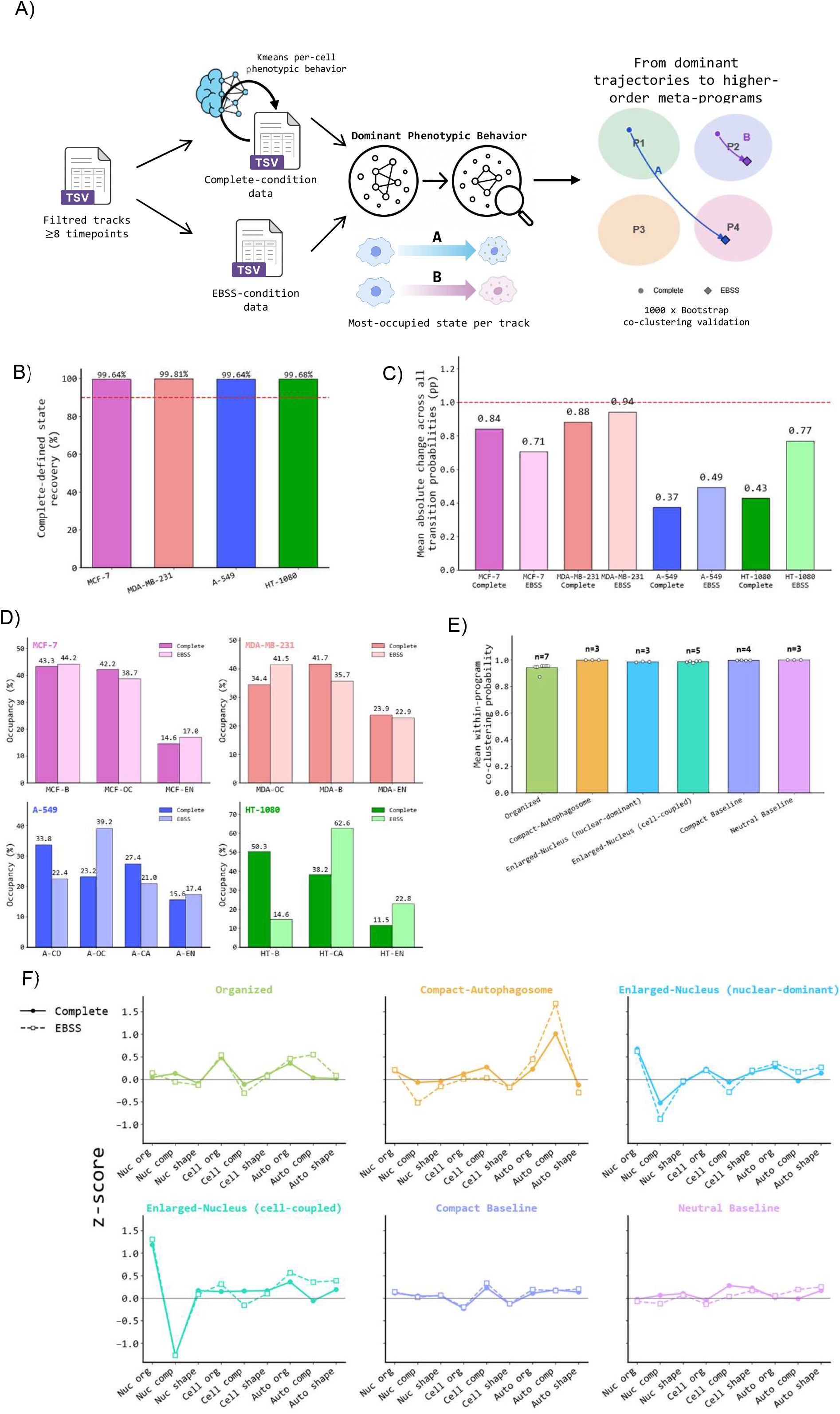
Robustness and program-assignment diagnostics for the state model. (A) Schematic of the program-assignment pipeline. Filtered tracks (≥ 8 timepoints) are split by condition: Complete-condition cells are K-means-clustered into per-cell phenotypic states, and EBSS-condition cells are projected into the same state space (Figure 5 / Methods). For each track, the most-occupied state across its own timepoints defines its Dominant Phenotypic Behavior. State pairs (within and across cell lines) are then tested for co-clustering across 1,000 bootstrap resamples of their nine-feature morphological signature; state pairs that consistently co-cluster are merged into a common program (schematic at right; P1–P4 and trajectories A/B are illustrative only, not real program identities). (B) State-recovery accuracy: percentage of Complete-defined state assignments correctly recovered after projecting EBSS cells into the same space, one bar per cell line (values labeled on each bar). Dashed red line: 90% reference threshold. (C) Bootstrap robustness of the Markov transition-probability matrix (1,000 resamples with replacement per cell line × condition); bars show the mean absolute change across the full transition matrix (values labeled on each bar). (D) Predicted state-occupancy distributions (% of cell-timepoints per state, Complete vs. EBSS), one panel per cell line. (E) Co-clustering membership confidence for each of the six trajectory programs (mean pairwise co-clustering probability among program members; 1,000-resample bootstrap consensus). (F) Weighted meta-feature signature profiles of the six trajectory programs, weighted by state occupancy (n cell-timepoints).

## AURORA: Supplementary Methods

### Detailed imaging acquisition settings

Time-lapse high-content imaging was performed on an Operetta system using a 20× long working distance objective. Before staining, residual growth medium was removed and cells were washed three times with PBS. Complete medium or EBSS was then added and cells were allowed to equilibrate for 15 min. Hoechst 33342 (0.1 µM), DAPGreen (2.5 µM; Dojindo, D676), and CellMask Deep Red (0.25 µM) were then added without additional wash steps, and cells were incubated for a further 45 min before imaging. DAPGreen is a small-molecule live-cell probe reported to label autophagic structures without transfection, to remain fluorescent across the autophagic process, and to correlate strongly with LC3-positive puncta [28]. The first acquired image was defined as timepoint 1 of the imaging sequence; cells were therefore exposed to the corresponding condition during the 60 min pre-imaging incubation. Images were acquired every 20 min for 4 h, collecting brightfield and fluorescence channels corresponding to Hoechst 33342, DAPGreen, and CellMask Deep Red. No z-stacks were acquired. Imaging was performed at empirically optimized fixed instrument heights for each channel throughout the time-lapse acquisition. Acquisition settings were as follows: brightfield, 40 ms exposure, transmitted light, instrument height 1.0 µm; Hoechst, 20 ms exposure, excitation 360-400 nm, emission 410-530 nm, instrument height 1.5 µm; DAPGreen, 200 ms exposure, excitation 460-490 nm, emission 500-550 nm, instrument height 3.5 µm; and CellMask Deep Red, 200 ms exposure, excitation 620-640 nm, emission 650-760 nm, instrument height 2.0 µm.

### Image processing for visualization

Raw images used for representative figures and visualization were processed in ImageJ/Fiji version 1.54p (Java 1.8.0_322, 32-bit) [36,37]. Channels were handled independently and converted to 16-bit format, followed by background subtraction. Contrast and threshold levels were adjusted only to improve visualization of Hoechst 33342, DAPGreen, and CellMask Deep Red signals. Composite images were generated for figure display. These ImageJ/Fiji processing steps were used exclusively for visualization and did not affect quantitative measurements used for downstream analyses.

### Detailed segmentation and feature-extraction workflow

Quantitative image analysis was performed in Harmony 4.8 using a compartment-aware segmentation workflow. Nuclei were segmented from the Hoechst channel and whole-cell regions from the CellMask Deep Red channel. Cytoplasmic masks were generated by subtracting the nuclear mask from the whole-cell mask. DAPGreen signal was background-corrected and used for object-level detection of DAPGreen-positive autophagosome-associated structures. For each segmented object or compartment, Harmony-derived morphological, morphometric, texture, and fluorescence intensity features were extracted. Segmentation thresholds were optimized separately for each cell line to account for cell-line-specific differences in morphology and signal intensity, but the same thresholds were applied to Complete and EBSS conditions within a given cell line. Objects touching image borders were removed using the “Remove Border Objects” function.

### Harmony analysis sequence

Segmentation and measurement extraction were performed using a standardized Harmony 4.8 analysis sequence. Input images were processed as individual planes, with brightfield correction enabled, no flatfield correction, no global image creation, and no Quick Tune step. Nuclei were identified in the DAPI/Hoechst channel using Harmony method A with a common threshold range of 0.20-0.60 and a minimum nuclear area of 30 µm². Border-touching nuclei were removed using common filters, generating the final selected nuclear population.

Whole-cell regions were segmented from the far-red CellMask channel using the selected nuclei as seeds. Cytoplasm detection used Harmony method B, with a common threshold range of 0.40-0.60 and individual threshold range of 0.60-1.00. Border-touching cell objects were removed, generating the final selected cell population. Intracellular cytoplasmic regions were defined from the selected cell cytoplasm using a common threshold of 0.00, split into objects, with a minimum area threshold of >0 µm² and without hole filling.

Cell morphology was first quantified on the selected cell population using standard Harmony morphology measurements, including cell area and roundness. A two-class linear classifier was then trained using cell area and cell roundness to separate countable cells from rounded or non-countable cells. The resulting Countable Cell population was used for downstream quantification.

DAPGreen-positive autophagosome-associated structures were detected in the fluorescein channel within the cytoplasm of Countable Cells using a common threshold range of 0.20-0.60, object splitting enabled, minimum area >0 µm², and no hole filling. Surrounding cell regions were defined in the far-red channel from Countable Cells using Harmony method A, with individual threshold range 0.20-0.40 and excluding the input cell region. This surrounding region was used for local fluorescence background estimation.

Standard morphology properties were extracted from Countable Cells for the nucleus, cytoplasm, and autophagosome-associated regions, including area, roundness, width, length, and width-to-length ratio where applicable. STAR morphology properties were additionally calculated for nuclear and autophagosome-associated regions, including symmetry, threshold compactness, axial, and radial descriptors. Standard intensity measurements were extracted as mean fluorescence intensity from the DAPI/Hoechst channel in the nucleus and from the fluorescein channel in cytoplasm, surrounding cell regions, and autophagosome-associated regions.

Object-level results were exported for the Countable Cell population, including object number and mean measurements. Corrected cytoplasmic fluorescein intensity was computed in Harmony as the difference between the mean cytoplasmic fluorescein intensity and the mean surrounding-region fluorescein intensity. Exported object-level tables were then used for track organization, feature engineering, quality control, and downstream modeling.

### Tracking and longitudinal organization

Harmony-derived object tracking was used as the source of longitudinal identity. Exported object-level measurements were organized into track-level tables across cell lines, conditions, biological replicates, wells, fields, and timepoints while retaining the corresponding compartment-resolved features. These tracked cells were then used for continuity filtering, displacement QC, state assignment, transition analysis, and downstream state-occupancy summaries.

### Minimum consecutive track-length filtering

Track continuity was quantified as the longest uninterrupted sequence of valid measurements (Lmax) for each tracked cell across the 13 timepoints acquired during the 4 h experiment. Retention curves were generated by computing the percentage of tracks retained across increasing minimum consecutive track-length thresholds (k = 6-13; Supplementary Fig. 1A). In the global pooled analysis, k = 9 was identified as the knee point, retaining 63.8% of tracks. However, k = 8 retained 67.8% of tracks globally and avoided excessive loss of analyzable tracks in conditions with lower track stability, particularly HT-1080 under EBSS, where retention decreased from 25.3% at k = 8 to 20.9% at k = 9. Therefore, k = 8 was selected as the final threshold to balance temporal continuity with retention of biologically informative longitudinal single-cell tracks. Tracks with Lmax < 8 were excluded from downstream analyses. This filtering reduced the contribution of partial or unstable tracks caused by field-of-view exit, border exclusion, intermittent segmentation, or tracking discontinuities.

### Definition of Total Autophagy

Autophagy-associated signal was quantified using DAPGreen-positive object intensity and area after threshold-based background correction. Let CIF denote the mean fluorescein intensity within the cytoplasmic or whole-cell region, and let CSIF denote the local surrounding-region fluorescein intensity used as a background reference. The corrected intracellular background reference was computed as:

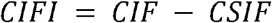

Let AIF denote the mean fluorescence intensity measured within DAPGreen-positive objects. Corrected autophagy intensity was computed as:

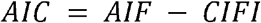

Total Autophagy (AT) was then defined as:

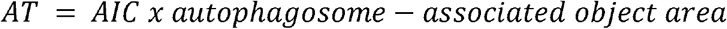

where autophagosome-associated object area corresponds to the summed area of detected DAPGreen-positive objects per cell, expressed in µm². Under this definition, AIC represents a corrected intensity term in arbitrary units, and multiplication by area yields a quantity proportional to integrated corrected fluorescence. This metric therefore captures both autophagosome-associated signal intensity and detected autophagosome-associated area per cell. Negative AIC values indicate that the measured object-level DAPGreen signal did not exceed the corrected intracellular background estimate under the segmentation and correction model used here. Because area-weighting such background-dominated values would generate sign-inverted integrated scores that are not interpretable as autophagosome-associated signal load, negative AIC values and the corresponding derived negative AT values were excluded from downstream analyses. This exclusion was intended to avoid misinterpreting background-dominated measurements as biologically meaningful “negative autophagy” values. The proportion of excluded negative AIC or AT measurements was monitored across cell lines and conditions as part of quality control. Background correction parameters were defined separately for each cell line and applied identically across Complete and EBSS conditions.

### Engineered morphology and organization features

Selected engineered features were computed in Python from Harmony-exported descriptors after segmentation, tracking, and longitudinal quality control. Features were calculated at the single-cell and timepoint level and, where applicable, separately for nuclear, cytoplasmic, whole-cell, and DAPGreen-positive autophagosome-associated compartments.

### Feret-based size approximation

To estimate object size efficiently, a simplified Feret-based descriptor was computed by taking the maximum value between each object’s measured length and width. This value was used as an approximation of the object’s largest structural dimension and served as an input for downstream shape-related descriptors.

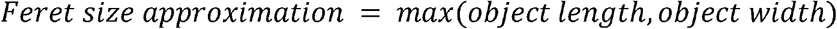

### Mass distribution

Mass distribution was defined as the proportion of a subcellular compartment area relative to the total cell area:

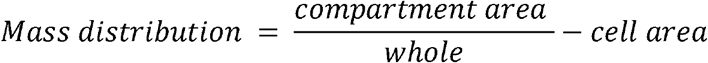

This feature provides a normalized descriptor of compartmental organization and was used to compare the relative spatial contribution of nuclear, cytoplasmic, and autophagosome-associated compartments across cells.

### Eccentricity

Eccentricity was used to quantify object elongation based on the relationship between the major and minor axes of an ellipse approximating the object. The semi-major axis a and semi-minor axis b were derived from the longest and shortest object diameters, respectively:

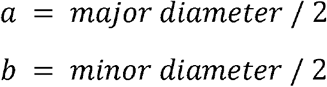

Eccentricity was calculated as:

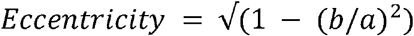

Values closer to 0 indicate more circular objects, whereas values closer to 1 indicate more elongated objects.

### Morphological irregularity index

A composite irregularity index was calculated to capture deviations from compact, circular morphology by integrating Feret-based size, circularity, minimum Feret diameter, and area-related descriptors. This metric was used as a computational proxy for combined elongation and shape irregularity.

### Fractality-related cell complexity descriptor

A fractality-related descriptor was computed as a proxy for cell size-associated structural complexity using a log-scaled area-to-reference-unit relationship:

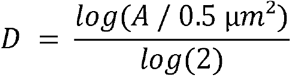

where A is the cell area in µm² and 0.5 µm² is the reference unit area. This descriptor was used as an approximate complexity-related feature rather than a true box-counting fractal dimension.

### pycytominer feature selection

Feature selection was performed using pycytominer [29] to remove non-informative and redundant variables before downstream analyses. Features were filtered using pycytominer-style feature-selection steps for missingness, near-zero variance, and pairwise correlation pruning. The same filtering logic was applied independently to each cell line, and pairwise correlation pruning used a correlation threshold of 0.95. The resulting cell-line-specific selected feature sets were used for subsequent phenotypic state modeling and state-occupancy analyses. Starting from the shared 83-feature universe, pycytominer retained 65 features for MCF-7, 65 for MDA-MB-231, 58 for A-549, and 56 for HT-1080 in the Complete-trained state models. The complete mapping between the shared descriptor panel, pycytominer-selected feature sets, and meta-feature groups is provided in a separate supplementary table.

### Meta-feature construction for landscape and state-occupancy program interpretation

For visualization and biological interpretation of phenotypic states and state-occupancy programs, image-derived descriptors were summarized into compartment-level meta-features. This analysis was performed independently of pycytominer feature selection. Whereas pycytominer-selected features were used for cell-line-specific phenotypic state model training, meta-features were used to interpret the resulting state landscapes and state-occupancy programs.

Meta-features were constructed by grouping shared image-derived descriptors into three compartments (nucleus, whole cell, and autophagosome-associated objects) and three feature axes (organization, compactness, and shape), yielding nine meta-feature groups: nuclear organization, nuclear compactness, nuclear shape, cell organization, cell compactness, cell shape, autophagosome organization, autophagosome compactness, and autophagosome shape. For each cell line, descriptors were z-normalized across Complete and EBSS cells before meta-feature aggregation. Meta-feature values were then computed as the mean of the corresponding grouped descriptors for each phenotypic state and condition.

The final landscape radar plots were generated from the full set of 83 shared image-derived descriptors present across all four cell lines. Each descriptor was assigned to one of the nine compartment-axis groups. Feret size approximation Nuclei was assigned to nuclear shape, Feret size approximation Autophagy to autophagosome shape, and Total Autophagy to autophagosome organization as an autophagosome-associated signal/load descriptor. The complete mapping between the original feature set, pycytominer-selected features for each cell line, and landscape meta-feature groups is provided in Supplementary Table 1.

### Complete-trained phenotypic state model

Additional details for the Complete-trained phenotypic state modeling workflow are provided below.

Phenotypic state modeling was performed independently for each cell line using the robust-normalized longitudinal single-cell dataset after quality-control filtering. The normalized matrices preserved the same filtered cell-time observations as the cleaned longitudinal datasets and therefore only included tracks with at least eight valid timepoints. Robust normalization was anchored to the Complete condition within matched replicate-timepoint strata. For each feature, cell values were centered relative to the median of the corresponding Complete-condition population from the same replicate and timepoint and scaled by the corresponding interquartile range. When the interquartile range was zero, a center-only transformation was applied.

Complete-condition cells were used to define the reference phenotypic state landscape for each cell line. Candidate K-means state structures were evaluated in the pycytominer-selected, robust-normalized feature space, whereas low-dimensional embeddings were used for visualization and projection of the resulting reference states. Candidate K values were ranked using silhouette and stability metrics, after which the selected solutions yielded three states for MCF-7, three for MDA-MB-231, four for A-549, and three for HT-1080. Final K-means fitting used random seed 42 and multiple initializations for reproducibility. The resulting Complete-defined state labels were treated as operational phenotypic reference states and were then used as supervised targets for cell-line-specific state classifiers.

For each cell line, the Complete-trained state model was implemented in scikit-learn as a multinomial logistic regression classifier with elastic-net regularization and the saga solver. Feature selection was first performed with pycytominer as described above, yielding a cell-line-specific modeling feature set. The classifier pipeline included constant-value imputation for missing measurements, feature standardization, and class-balanced multinomial fitting (class_weight = balanced). Hyperparameters controlling regularization strength (C) and elastic-net mixing (l1_ratio) were selected by stratified cross-validation within the Complete-condition dataset using negative log loss as the scoring criterion. Final archived model objects used max_iter = 3000, tol = 1e-4, and random_state = 0. The selected hyperparameters were C = 4.6416 and l1_ratio = 0.5 for MCF-7 and MDA-MB-231, and C = 1.0 and l1_ratio = 0.9 for A-549 and HT-1080. To limit overrepresentation of abundant states during model fitting, Complete-condition training rows were downsampled per state when necessary while preserving state balance.

After model fitting, the Complete-trained classifier was applied unchanged to EBSS-condition cells from the same cell line. This projection step assigned EBSS cells to the Complete-defined reference states and yielded per-cell predicted states and class probabilities in a common phenotypic state framework. The same model was also applied back to Complete cells for recovery and internal consistency analyses.

### Markov transition analysis

Markov transition analyses were computed separately for each cell line and condition using ordered single-cell state sequences. Consecutive timepoints were used to tabulate state-to-state transitions for every tracked cell. Transition probabilities were calculated by normalizing each row of the transition count matrix by the total number of outgoing transitions from the corresponding state, thereby retaining both inter-state switching and self-transition stability. Dominant outgoing transitions and state stability probabilities were used for visualization and summary.

Net displacement was calculated as the Euclidean distance between the first and last valid Harmony-exported object centroid coordinates for each retained track and was used only as a longitudinal QC summary.

### Trajectory-program discovery via bootstrap co-clustering

Dominant trajectories were derived from the full longitudinal state assignments of individual tracked cells (see Complete-trained phenotypic state model, above) and used to build per-group centroids that summarize longitudinal state usage together with the associated meta-feature signature. For each retained track, predicted states across the longitudinal sequence were converted into a state-occupancy vector, defined as the fraction of measured timepoints assigned to each reference state; the dominant trajectory for that track corresponded to the state with the highest occupancy across its retained valid timepoints. Tracks were grouped by cell line, condition, and dominant trajectory into 26 CellLine × Condition × State groups. For each group, a centroid was constructed as the median of the nine compartment-level meta-features (see Meta-feature construction, above) across that group’s member tracks.

Rather than relying on a single UMAP/HDBSCAN run on these 26 group centroids — which is sensitive to the specific random draw of tracks contributing to each centroid — group similarity was instead assessed through a bootstrap co-clustering procedure (below), and a final, deterministic program assignment was obtained from the resulting co-clustering probabilities rather than from any single embedding.

### Bootstrap co-clustering and program-assignment parameters

The bootstrap procedure operated on the same 26 CellLine × Condition × State groups described above, using each group’s own set of per-track, robust-normalized nine-dimensional meta-feature vectors (one row per track, already summarized as that track’s own median across its retained timepoints) as the resampling unit.

In each of 1,000 bootstrap iterations, tracks were resampled with replacement independently within each of the 26 groups, and each group’s centroid was recomputed as the median of the resampled tracks’ meta-feature vectors, yielding a new 26×9 centroid matrix. This matrix was standardized (z-scored per column) and embedded into two dimensions with UMAP (Euclidean metric, n_neighbors = 6, min_dist = 0.20, random_state set to the iteration index for reproducibility). HDBSCAN (Euclidean metric, min_cluster_size = 3, min_samples = 2, cluster_selection_method = eom) was then applied to the resulting embedding. For every pair of the 26 groups, a co-clustering counter was incremented whenever both groups were assigned to the same non-noise HDBSCAN cluster in that iteration.

After all iterations, the co-clustering probability for each pair of groups was defined as its co-clustering count divided by the number of completed iterations (1,000 of 1,000 requested). A final, deterministic program assignment was then obtained by applying HDBSCAN with the same parameters (Euclidean metric, min_cluster_size = 3, min_samples = 2, cluster_selection_method = eom, no random seed) to the 26×26 distance matrix defined as one minus this co-clustering probability.

This procedure identified six trajectory programs, each internally supported by co-clustering probabilities among its member groups (mean within-program co-clustering probabilities summarized in Supplementary Fig. 4E), plus one residual group — a single HT-1080/EBSS state — that did not reach reproducible co-clustering with any other group and was retained separately as unassigned (“No program”). This assignment was stable when the bootstrap was independently repeated at 300 and 1,000 iterations: the same six programs and the same unassigned group were recovered in both runs, with only one borderline group changing which program it co-clustered with most strongly between the two runs, consistent with expected refinement from additional resampling.

For visualization only (Fig. 5D), the same 26 group centroids (nine meta-features, computed from the observed data without resampling) were projected into a shared two-dimensional space using principal component analysis, fit once across all four cell lines. Each cell-line panel shows this same shared projection and the same program regions, cropped to the region occupied by that cell line’s own centroids.

## Notes

### Competing Interest Statement

The authors have declared no competing interest.

### Summary of Updates

This version represents a substantial revision and expansion of the manuscript. The study has been redesigned as a comprehensive framework for the dynamic tracking and single-cell analysis of autophagic responses. New cell lines and experimental models have been incorporated, together with additional experiments and analyses. The methodology, results, figures, and supplementary materials have been extensively revised and expanded to reflect the new scope of the study. The manuscript has also been reorganized to better present the framework and its applicability for the analysis of autophagic responses at the single-cell leve

